# Transmission of *Klebsiella* strains and plasmids within and between Grey-headed flying fox colonies

**DOI:** 10.1101/2021.10.25.465810

**Authors:** Ben Vezina, Louise M. Judd, Fiona K. McDougall, Wayne S.J. Boardman, Michelle L. Power, Jane Hawkey, Sylvain Brisse, Jonathan M. Monk, Kathryn E. Holt, Kelly L. Wyres

## Abstract

The Grey-headed flying fox (*Pteropus poliocephalus*) is an endemic Australian fruit bat, known to carry pathogens with zoonotic potential. We recently showed these bats harbour the bacterial pathogens *Klebsiella pneumoniae* and closely related species in the *K. pneumoniae* species complex (*Kp*SC). However, the dynamics of *Klebsiella* transmission and gene flow within flying fox colonies were not explored and remain poorly understood.

Here we report a high-resolution genomic comparison of 39 *Kp*SC isolates from Grey-headed flying foxes. Illumina whole genome sequences (n=39) were assembled *de novo* and the Kleborate genotyping tool was used to infer sequence types (STs). Oxford Nanopore sequences were generated for 13 isolates (one for each distinct ST) in order to generate high-quality completed reference genomes. Read mapping and variant calling was used to identify single nucleotide variants (SNVs) within each ST, using the relevant reference genome. *In silico* genome-scale metabolic models were generated to predict and compare substrate usage to 59 previously published *Kp*SC models for isolates from human and environmental sources, which indicated no distinction on the basis of metabolic capabilities.

High-resolution genome comparisons identified five putative strain transmission clusters (four intra- and one inter-colony, n=2-15 isolates each, ≤25 pairwise SNVs). Inter-colony transmission of *Klebsiella africana* was found between two flying fox populations located within flying distance. The 13 completed genomes harboured 11 plasmids, all of which showed 37-98% coverage (mean 73%) and ≥95% identity to those previously reported from human-associated *Kp*SC. Comparison of plasmids from different flying fox associated *Kp*SC indicated an interspecies horizontal plasmid transmission between *K. pneumoniae* and *K. africana* for a 98 kbp plasmid, pFF1003.

These data indicate that *Kp*SC are able to transmit directly via flying fox populations or indirectly via a common source, and that these isolates can harbour plasmids with similarity to those found in human derived *Kp*SC, indicating gene flow is occurring between isolates from Grey-headed flying fox *Kp*SC and human clinical isolates.

## Introduction

The *Klebsiella pneumoniae* species complex (*Kp*SC) [2] is a group of closely-related *Klebsiella* species which includes *Klebsiella pneumoniae, Klebsiella quasipneumoniae* subsp. *quasipneumoniae, Klebsiella variicola* subsp. *variicola, Klebsiella quasipneumoniae* subsp. *similipneumoniae, Klebsiella variicola* subsp. *tropica, Klebsiella quasivariicola* and *Klebsiella africana. Kp*SC are problematic as they function as cosmopolitan opportunistic pathogens [3, 4], responsible for a worrying proportion of community and hospital-acquired infections [5]. Prevalence of multi-drug resistance and acquired virulence factors associated with invasive infections is increasing over time [6], hence identifying the reservoirs of problematic *Kp*SC lineages/sequence types (STs), and mobile antimicrobial resistance and virulence determinants, is key for targeting interventions to limit the spread of these organisms.

Aside from humans, *Kp*SC have also been isolated from a wide variety of environments including soil [7], plants [3, 8] fresh water [7, 9], marine environments and organisms [10], waste dumps [11], animals including cats and dogs [12], migratory and domesticated birds [13, 14], agricultural animals [15, 16] and Grey-headed flying foxes [1]. Flying foxes, or fruit bats, have long been considered important vectors of zoonotic viruses such as Australian bat lyssavirus [17], Menangle virus [18] and Hendra virus (though not as the primary reservoir) [19]. There is growing evidence that Grey-headed flying foxes also harbour human pathogenic bacteria such as antibiotic resistant and pathogenic *Escherichia coli* lineages [20]. Carbapenem resistant *K. pneumoniae* have also previously been isolated from insectivorous bat guano in Algeria [21], though at very low frequency (two positive out of 110 samples).

Our previous surveillance study [1] identified *Kp*SC from three of four wild flying fox colonies (up to 36% prevalence per colony) and a captive rehabilitation colony (80% prevalence) in Australia (**Figure 1**). Notably, this was the first non-human sighting of *K. africana*, a species which has been reported in just two previous studies globally [6, 22]. Our initial genomic analyses of these *Kp*SC strains from flying foxes indicated low ST diversity with few STs previously reported among human clinical isolates, as well as low rates of antimicrobial resistance (1.1%) and virulence factors, suggesting that these Grey-headed flying fox colonies are not a high-risk environmental reservoir for problematic *Kp*SC isolates. However, this analysis did not explore detailed strain relationships, nor the broader genetic content of these *Kp*SC.

**Figure 1:**
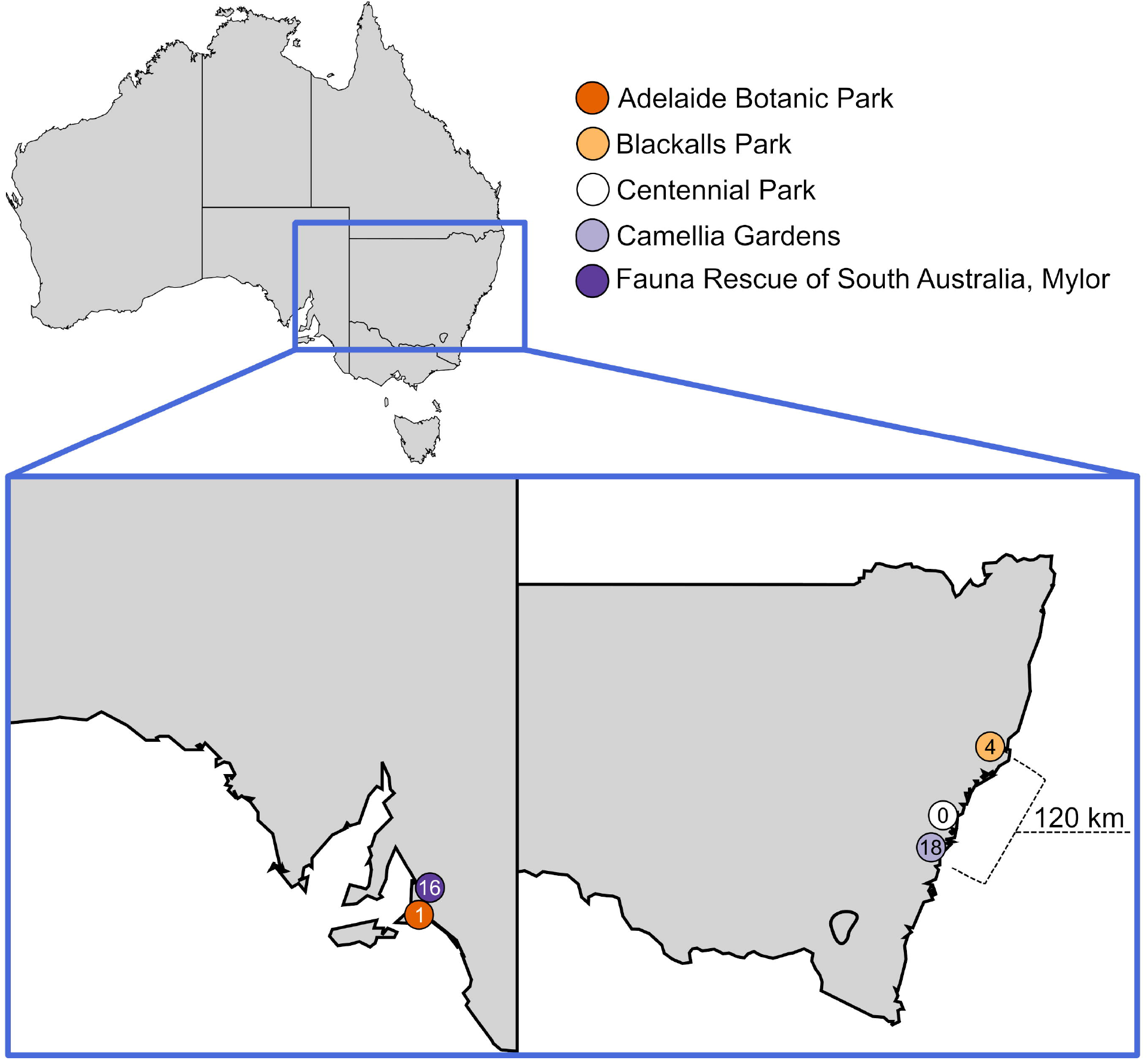
Sampled flying fox colonies across Australia by McDougall *et al* [1]. Coloured circles represent grey-headed flying fox colonies and the number within represents the number of *Klebsiella pneumoniae* species complex (*Kp*SC) isolates obtained. *Kp*SC were obtained from all colonies except for Centennial Park. Map of Australia generated by Lokal_Profil from https://upload.wikimedia.org/wikipedia/commons/b/bf/Australia_map%2C_States.svg, used as per the Creative Commons Attribution-Share Alike 2.5 Generic license.

Here, we use long-read DNA sequencing to complete the genomes of 13 *Kp*SC from flying foxes and leverage these data alongside the previously reported draft genomes to perform the first high-resolution analysis of *Kp*SC strain and plasmid transmission dynamics within/between Grey-headed flying fox colonies. To better understand the potential for strain and plasmid transfer into the human population, we compare completed plasmid sequences from flying fox isolates to those identified among human-associated *Kp*SC, and utilise the latest genome-scale metabolic modelling approaches to compare the metabolic capabilities of flying fox derived isolates to *Kp*SC isolates from other sources.

## Results

### High-resolution SNV analysis identified multiple putative strain transmissions

Among the 13 distinct *Kp*SC STs previously reported from our flying fox isolate collection, five were associated with >1 isolate each (**Figure 2**), four of which comprised pairs or groups of isolates that differed by ≤25 SNVs, indicative of recent strain transmissions [31–33]. Mapping data can be found in **Table S2**, while pairwise SNVs can be found in **Table S3**, ranging from 0-715.

**Figure 2:**
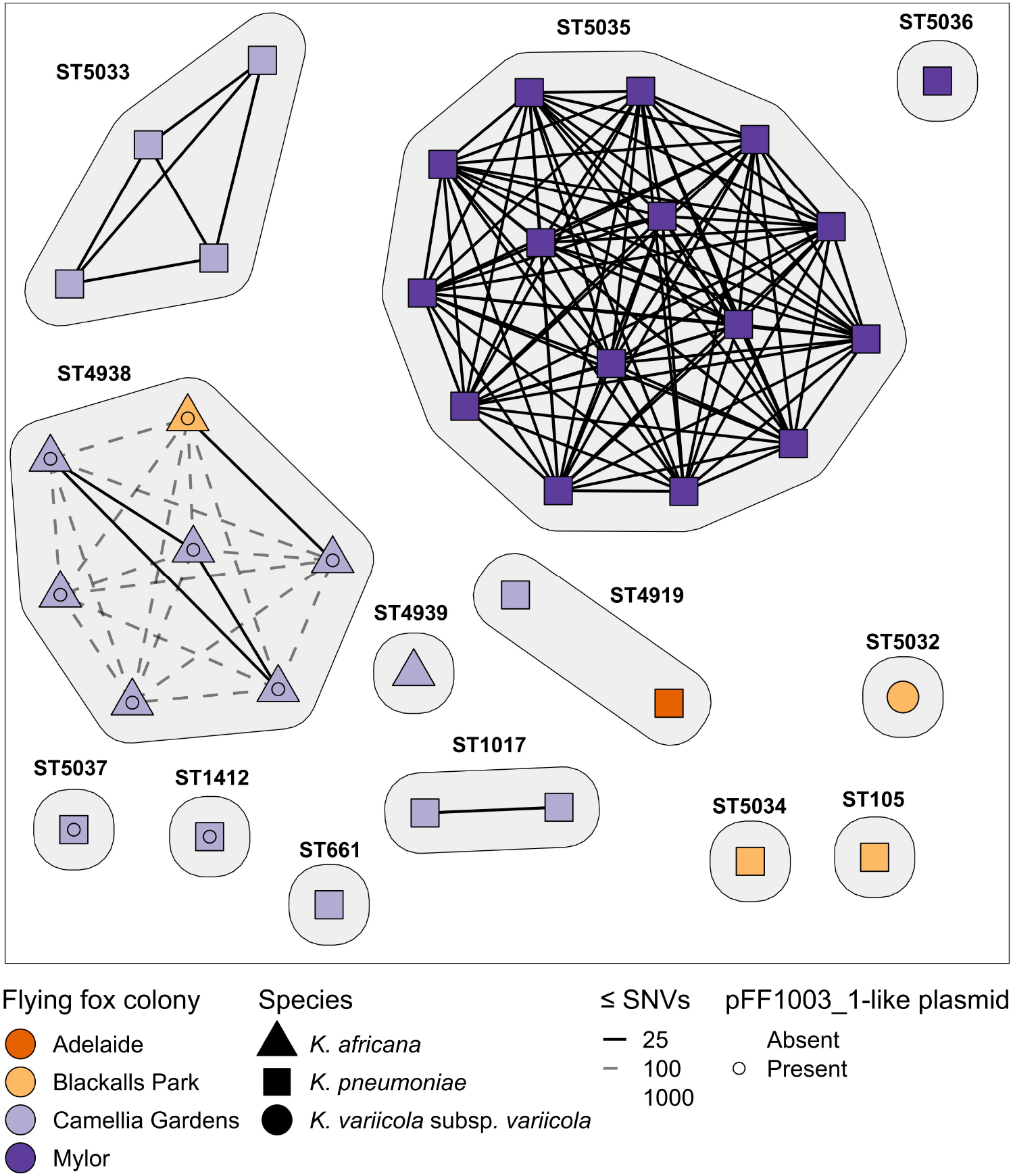
Network graph showing pairwise single nucleotide variant (SNV) distances between isolates of the same sequence type. Isolates are shown as nodes, grouped by sequence type (ST) with the shape representing the species. The SNV distances are shown as lines between nodes. Isolates are considered part of a transmission cluster if they have ≤25 pairwise SNVs, represented by a solid line. Dashed lines show 26 - 100 pairwise SNVs. Isolates are coloured based on their Grey-headed flying fox colony isolation source as indicated in the figure. The presence of the pFF1003-like plasmid is also indicated.

There were three occurrences of putative intra-colony transmission at the Camellia gardens colony, including *K. pneumoniae* ST1017 (n=2 isolates, three pairwise SNVs), *K. africana* ST4938 (n=3 isolates, zero pairwise SNVs) and *K. pneumoniae* ST5033 (n=4 isolates, ≤2 pairwise SNVs), and one occurrence at the captive Mylor colony of *K. pneumoniae* ST5035 (n=15 of 16 isolates recovered from this colony, ≤3 pairwise SNVs) (**Figure 2**). A single putative *K. africana* ST4938 inter-colony transmission event was observed between the Camellia Gardens and Blackalls Park colonies (n=2 isolates, 21 pairwise SNVs). Notably, this inter-colony cluster was separated from the Camellia Gardens ST4938 intra-colony cluster and two unclustered isolates by just 36-58 pairwise SNVs, which may be indicative of longer-term circulation within the colony and/or acquisition from a common long-term reservoir (also see ST4938 phylogeny below).

Two *K. pneumoniae* ST4919 isolates, one from the Adelaide colony and the other from the Camellia Gardens colony (approximately 1,150 kilometres apart), were separated by 715 SNVs.

### Plasmid diversity and transmission between species

A total of 11 plasmids were identified across the 13 completed genomes. Two isolates (*K. africana* ST4938, strain FF1003 and *K. pneumoniae* ST5033, strain FF979) harboured two plasmids each, seven isolates harboured a single plasmid each, while four isolates contained none (**Table S2**). The plasmids ranged from approximately 5 kilobases (kb) to 175 kb in length (**Figure 3, Table S4**). At least 7 of the 11 plasmids were predicted to be either self-conjugative or mobilizable and only one contained any antimicrobial resistance genes (*K. pneumoniae* ST1017, strain FF1009_1, carrying *qnrS1* and a variant of *dfrA14*) and none contained any of the well characterised *Kp*SC virulence genes [2]. Nonetheless, all plasmids demonstrated high identity (≥95%) with generally high coverage (37-98%; mean 73%) to at least one plasmid from human-derived *Kp*SC isolates (**Figure 3**).

**Figure 3:**
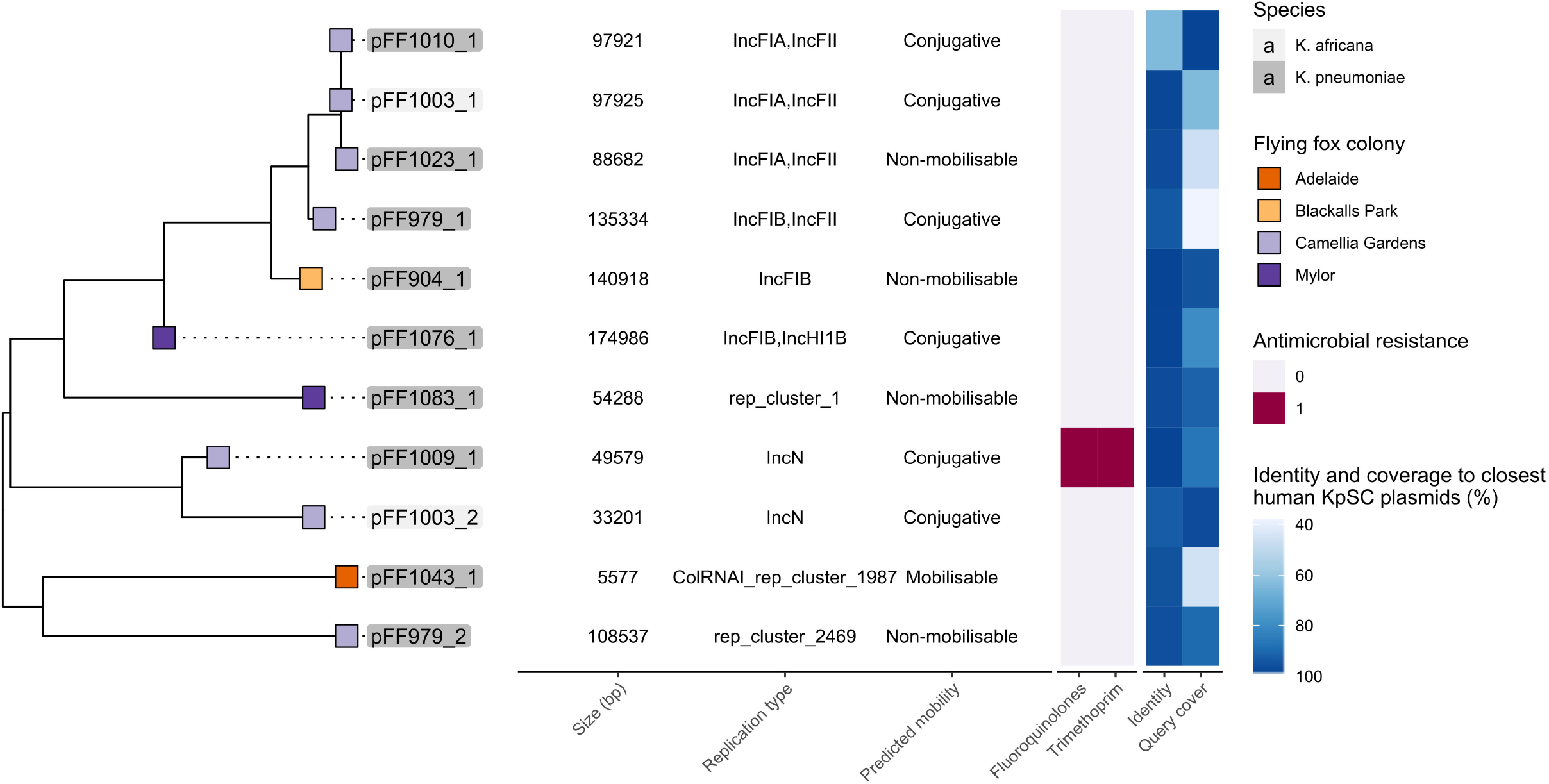
MASH distance tree of plasmids found in the long read flying fox *Kp*SC genomes. Tree was inferred from MASH distances using the FastME algorithm. Tip colour indicates grey-headed flying fox colony, while colour of plasmid label denotes the species as per legend in figure. The corresponding heatmap shows presence of antimicrobial resistance genes in pink, along with identity and coverage to the closest *Kp*SC plasmid isolated from a human source, identified by Genbank’s BLASTn. Accession numbers can be found in **Table S4**.

A reference-based mapping approach was used to determine the presence or absence of each of our completed plasmids among host isolates of the same ST, which revealed a high level of conservation ≥78% mapping coverage in all but three instances, comprised of three out of 7 *K. africana* ST4938 isolates missing the pFF1003_2 plasmid) (**Table S2**). At most, plasmids differed amongst themselves by eight pairwise SNVs (**Table S3**). Notably, these included the pFF1009_1 AMR plasmid identified in both *K. pneumoniae* ST1017 isolates, and the ~5.5 kb pFF1043_1 plasmid identified among the *K. pneumoniae* ST4919 isolates collected from two independent colonies (Adelaide and Camelia Gardens). These contained zero and one pairwise SNVs to the completed pFF1009_1 and pFF1043_1, respectively.

Read mapping analyses indicated that a pFF1003_1-like conjugative plasmid was present in nine of the total 39 isolates (23%) (including those from both the Camellia Gardens and Blackalls Park colonies, maximum 8 pairwise SNVs). These included representatives of *K. africana* ST4938, as well as *K. pneumoniae* ST5037 and ST1412 (**Figure 3, Figure 4**). Comparison of the completed sequences for pFF1003_1 (mapping reference from FF1003, a *K. africana* ST4938 isolate) and pFF1010_1 (from FF1010, a *K. pneumoniae* ST1412) showed 99% BLASTn coverage and 99% identity, indicative of recent plasmid transmission, though SNV distances were not calculated. However, a comparison between the pFF1003_1 reference and the completed sequence for pFF1023_1 (from FF1023, a *K. pneumoniae* ST5037) showed only 77% BLASTn coverage and 99% identity. Approximately 9 kb of pFF1023_1 was deleted compared to pFF1003_1 (at least six independent deletion events), including part of the conjugation locus. As such, this plasmid was predicted to be non-conjugative. The pFF1023_1 *nan* locus (consisting of *nanA, nanT, nanE2, nanK, yhcH*, a putative N-acetylneuraminic acid porin (Genbank accession: HAT1683918.1), *nanC* and *nanM*, flanked by IS3 family transposases), was also inverted compared to the pFF1003_1 reference plasmid (**Figure 4C**).

**Figure 4:**
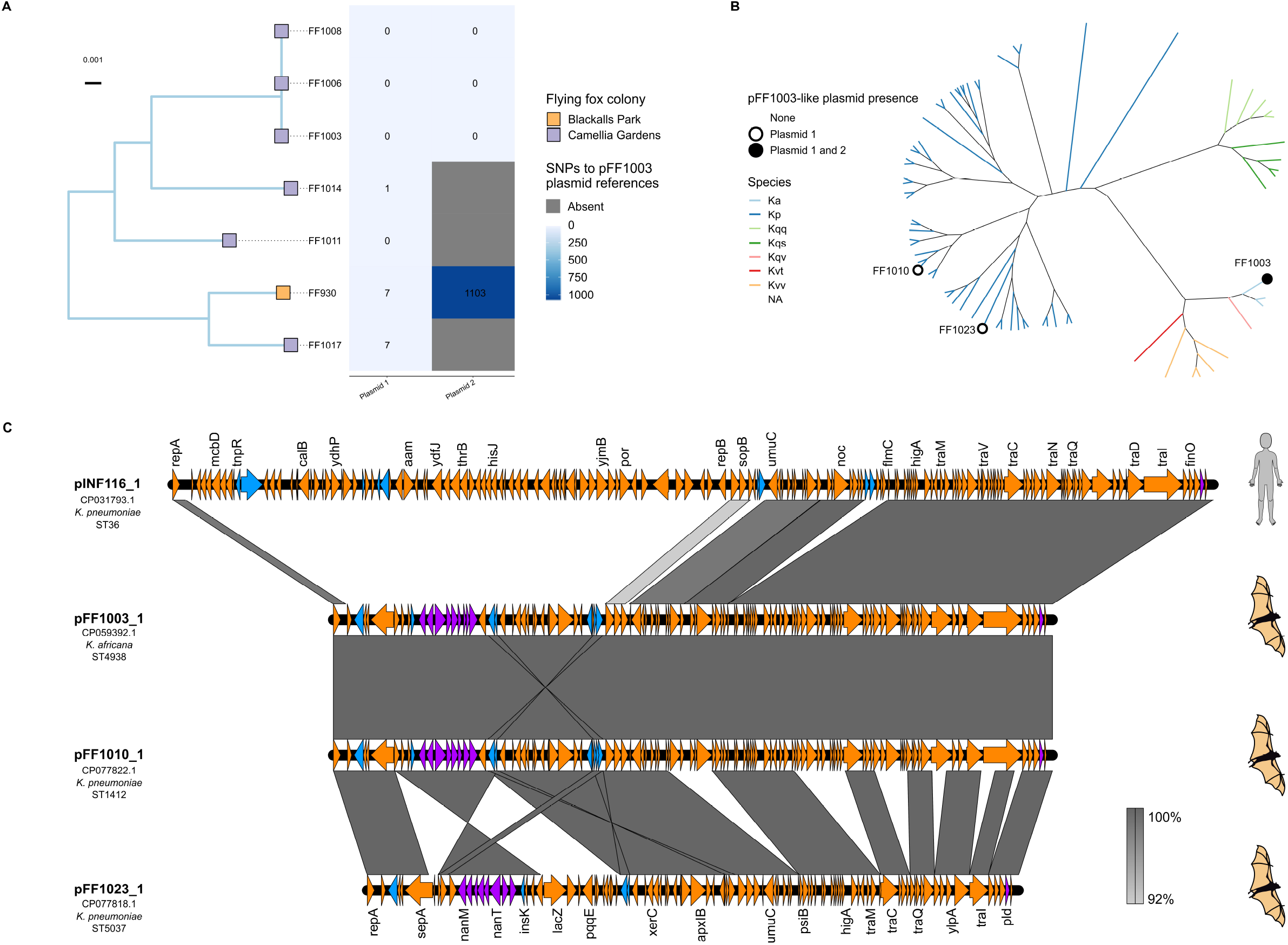
Comparison and distribution of ST4938 strains and associated plasmid FF1003_1. **A** Mid-point rooted core SNV chromosomal maximum-likelihood phylogeny of seven *K. africana* ST4938 isolates. Colony is indicted by tip colour. The heatmap shows plasmid SNV counts compared to the pFF1003_1 and pFF1003_2 completed reference plasmids. **B** Unrooted core chromosomal maximum likelihood daylight phylogeny of all flying-fox derived isolates included in the study with dots showing presence of pFF1003-like plasmids. Branch lengths were left unscaled to fit figure proportions. **C** Plasmid comparisons between pINF116_1 carried by *K. pneumoniae* INF116, isolated from a human source and flying fox plasmids pFF1003_1, pFF1010_1 and pFF1023_1. The orange arrows indicate genes, blue arrows indicate transposons and purple arrows indicate metabolic genes such as the sialic acid catabolism *nan* locus. The grey bars between maps indicate nucleotide identity as per legend in the figure.

pFF1003_1 (97 kb) showed 64% BLASTn coverage and 99% identity to the publicly available 141 kb human-derived *Kp*SC plasmid (pINF116_1). This plasmid was identified from *K. pneumoniae* ST36 strain INF116, isolated from an Australian human urine sample in 2013 (Genbank accession: CP031793.1). Compared with pINF116, pFF1003_1 lacked many of the amino acid transport genes such as *hisQ*, *hisJ* and *glnQ*, but contained the additional tyrosine recombinase *xerC*, tRNA(fMet)-specific endonuclease *vapC* and the putatively intact sialic acid degradation *nan* locus (**Figure 4C**).

### *Kp*SC from flying foxes were not distinguished by metabolic capabilities

We simulated growth phenotypes *in silico* and overlaid these on the core genome phylogeny of 72 *Kp*SC genomes to identify any potential genetic or metabolic commonalities of *Kp*SC isolates from Grey-headed flying foxes compared to those from other sources. All *Kp*SC isolates were predicted to utilise a core set of substrates accounting for 49.6% of carbon, 40% nitrogen, 72.9% of phosphorus and 40% of sulphur substrates tested (**Table S5, Table S6**). Percentage of total nutrient sources tested which displayed variable usage (non-core) across the *Kp*SC isolates were much lower, consisting of 10.7% carbon, 3.2% nitrogen, 1.7% phosphorus and 0% sulphur sources.

We generated a core gene phylogeny and overlaid the variable substrate usage profiles to examine the relationship between lineage and metabolic substrate usage (**Figure 5, Figure S1**). Isolates from flying foxes were distributed throughout the phylogeny and predicted growth phenotypes appeared to cluster by species, then more granularly by ST (**Figure 5**), rather than isolate source. Hierarchical clustering of isolates based on predicted growth phenotypes was also consistent with this observation (**Figure S2**), with isolates of the same species generally clustering together. One isolate, *K. pneumoniae* DHQP1002001 appeared as an outgroup to the other *K. pneumoniae* isolates in the phylogeny. This appeared to be a genetic species hybrid of mostly *K. pneumoniae* with roughly 1 Mb from a *K. quasipneumoniae* subsp. *quasipneumoniae* strain (**Figure S3**). Notably, unlike the rest of the *K. pneumoniae* isolates, this species hybrid was predicted to be unable to utilise ethanolamine and D-glucarate as sole sources of carbon. While *K. quasipneumoniae* subsp. *quasipneumoniae* isolates were universally predicted to be unable to utilise the ethanolamine, they were all predicted to utilise tricarballylate and D-glucarate. It is unclear whether this pattern of substrate usage loss exists for *Klebsiella* hybrid species in general or was limited due to the data used.

**Figure 5:**
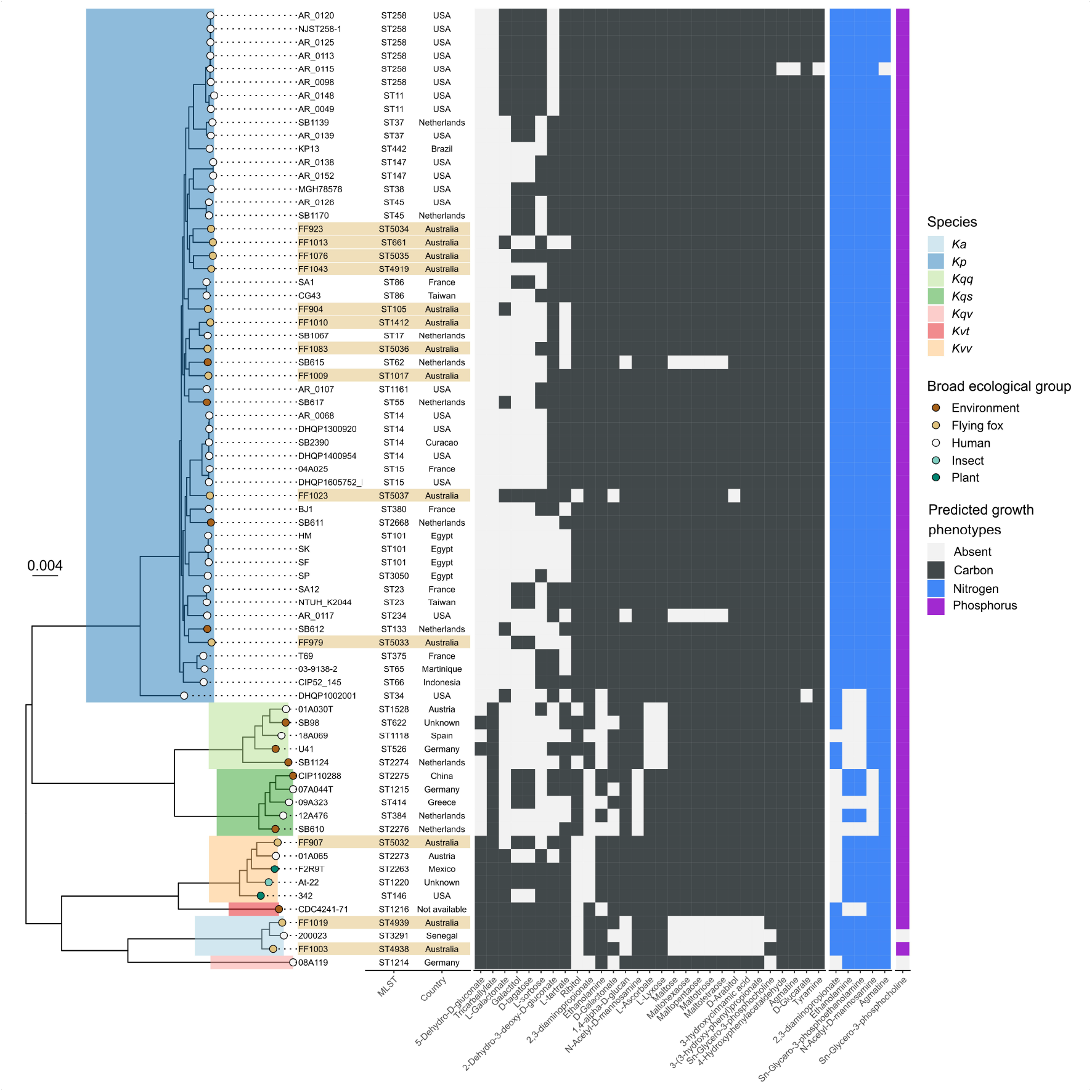
Mid-point rooted core genome maximum likelihood phylogeny of 72 KpSC isolates and variable metabolic substrate use. Species are shaded on the phylogeny branches as per legend in the figure; *Ka* refers to *K. africana, Kp* refers to *K. pneumoniae, Kqq* refers to *K. quasipneumoniae* subsp. *quasipneumoniae, Kqs* refers to *K. quasipneumoniae* subsp. *similipneumoniae, Kqv* refers to *K. quasivariicola, Kvt* refers to *K. variicola* subsp. *tropica, Kvv* refers to *K. variicola* subsp. *variicola*. Broad ecological group/ isolation source is shown by tip colour. Flying fox isolates shown as light brown strips. Bootstrap values are shown in **Figure S1**. Variable predicted growth substrate utilisation is shown in the heatmap, coloured by substrate type as indicated. No variable sulphur usage was predicted.

## Discussion

Our high-resolution genomic comparison of *Kp*SC from Grey-headed flying fox colonies across Australia indicates strong evidence for strain transmission both within and between colonies (**Figure 2**). Five putative strain transmission clusters were identified, four of which involved the wild Camellia Gardens colony in the state of New South Wales. This included three clusters contained within this colony and one transmission cluster between this colony and the wild colony at Blackalls Park. The latter was associated with *K. africana* ST4938 for which isolates FF1017 (Camelia Gardens) and FF930 (Blackalls Park), were separated by just 21 chromosomal SNVs (**Figure 2**) and shared a pFF1003_1-like (~98 kb) plasmid with 0 pairwise SNVs, indicating they descended from a very recent common ancestor (although a second plasmid present in FF1017 was absent from FF930). Notably, these colonies are situated 123 km apart (**Figure 1**) and were both sampled within a 23-day period. As Grey-headed flying foxes have been shown to travel large distances of ~50 km [67] and up to >1,000 km in some cases [68], it is possible for individual flying foxes to move between the Blackalls Park and Camellia Gardens colonies causing direct inter-colony transmission. Alternatively, transmission may have occurred indirectly via a common source (food/water etc). Additionally, our analyses indicated that the two ST4938 clusters, as well as two additional unclustered ST4938 isolates from Camellia Gardens differed by no more than 58 pairwise SNVs (**Figure 2, Table S3**). These values are much lower than would be expected for randomly selected *Kp*SC of the same ST [32, 69] and similar to those observed for isolates representing longer-term transmission within hospital networks [32, 69]. Hence, these findings are consistent with longer-term transmission in the flying fox population or acquisition from the same long-term reservoir.

The fifth putative strain transmission cluster was identified at the captive Mylor colony located within a rehabilitation centre, which at the time of sampling was a closed population of 30 – 40 adult Grey-headed flying foxes from the local wild Adelaide Botanic Park colony that had been affected by a heat stress event. These unhealthy individuals may have suffered increased susceptibility to opportunistic colonisation and dissemination of *Kp*SC isolates e.g. from other wildlife or domestic animals that were housed in close proximity, which we speculate facilitated the broad scale dissemination of *K. pneumoniae* ST5035 in the population (isolated from 15 of 20 sampled individuals [1]).

There is currently very limited data on the dynamics of bacterial transmission among flying foxes and other bat species; however, our results align with a previous study of viral dynamics [70], which found transmission of Nipah virus in Indian flying foxes within Bangladesh occurred within and between colonies. This study also noted the cycles of Nipah virus prevalence could be influenced by colony demographics, such as individual bat turnover which reduced herd immunity [70]. Grey-headed flying fox colonies in New South Wales have an average daily individual turnover of 17.5% as they travel between roosts via a node-based network [71], indicating high influx and efflux of individuals. This behavioural activity may make Grey-headed flying fox colonies susceptible to between-colony transmission of microbes such as *KpSC*, however further sampling is required to quantify these transmission events and determine if they represent transient or ongoing phenomena.

In addition to strain transmissions, we identified putative transmission of a 98 kb plasmid, passing the species barrier between *K. africana* ST4938 and *K. pneumoniae* ST1412 and ST5037 (**Figure 3, Figure 4**). The plasmid variants harboured by FF1003 (*K. africana* ST4938) and FF1010 (*K. pneumoniae* ST1412) were highly similar (4 pairwise SNVs, 100% coverage and 99% nucleotide identity), supporting a recent transmission event. In contrast, the plasmid variant harboured by FF1023 (*K. pneumoniae* ST5037) showed at least six deletions as well as a structural rearrangement, indicating a plasmid transmission likely occurred at some point in the more distant evolutionary history of these strains (either directly or indirectly via intermediary hosts). In that regard, it is possible that the pFF1003-like plasmid may have been circulating within the Camelia Gardens colony for some time and/or has been introduced from a common reservoir multiple times. Aside from the presence of several selfish genetic elements including a *higAB* toxin-antitoxin system, intact conjugative *tra* locus (except in the case of pFF1023_1) (**Figure 4C**), this pFF1103-like plasmid contained a putatively intact *nan* sialic acid locus that was conserved in all variants. The *nan* locus encodes degradation of N-acetylneuraminic acid, a sialic acid [72]. Utilisation and acquisition of N-acetylneuraminic acid improves colonisation and provides competitive growth advantages to a variety of pathogens [73–75]. It is possible that the presence of this locus provided some sort of selective advantage to promote the transmission and longer-term maintenance of the pFF1103-like plasmid in the Camelia Gardens population, although further investigations would be required to confirm this hypothesis.

Plasmid pFF1003 also showed high similarity to a plasmid identified from INF116, a *K. pneumoniae* clinical isolate causing a urinary tract infection in a patient in a Melbourne hospital in 2013 [76]. In fact, our search of publicly available nucleotide sequence data indicated that all completed plasmids from flying fox derived *Kp*SC shared high identity (≥95%) with generally high coverage (37-98%; mean 73%) (**Table S4**) to at least one other plasmid identified from human clinical isolates. Hence, our data provide clear evidence for the flow of genetic material either directly or indirectly between *Kp*SC isolated from flying foxes and human specimens, although we cannot speculate on the directionality or time-frame of transfer. In order for direct transfer to occur these isolate populations would need to coexist, at least transiently, within the same spatial-temporal and ecological niche.

While much more work is required to fully understand the distribution and flow of *Kp*SC strains and genetic material between niches (including detailed genomic analysis of large contemporaneous isolate collections), the results of the preliminary metabolic analyses presented here are consistent with niche overlap (i.e. *Kp*SC from flying foxes did not appear to be metabolically distinct). However, we note that our analysis was constrained to a small sample size and that there are likely many metabolic traits which are uncaptured within the metabolic models used in this study, due to limitations of the current input database. Over time, this approach should become more robust as databases become more complete (https://www.genome.jp/kegg/docs/upd_map.html) [77]. There may also be other genetic differences between flying fox and human strains of *Kp*SC such as K/O loci, previously discussed within the context of flying fox isolates by McDougall et al [1] however due to a paucity of Grey-headed flying fox isolates, we lack the statistical power to discern this.

## Experimental procedures

### Bacterial isolate collection and whole-genome sequencing

Thirty-nine *Kp*SC isolates including thirty *K. pneumoniae*, one *K. variicola* subsp. *variicola* and eight *K. africana* were previously isolated from four different Grey-headed flying fox colonies [1]; Blackalls Park, sampled on 12/12/2017 and 10/4/2018; Camellia Gardens, 4/1/2018; Adelaide Botanic Park, 11/2/2018, Mylor (captive colony), 12/2/2018; Centennial Park, 9/3/2018. One isolate (*K. pneumoniae* FF996) was not utilised in this study due to DNA contamination issues identified during high-resolution analysis. See **Table S1** for details of all isolates, genotyping results and genome accessions.

Illumina (short-read) whole-genome sequencing data was generated and reported previously [1]. DNA was extracted for long-read sequencing of 13 isolates representing distinct STs (see **Table S1**), using the GenFind v3 kit (Beckman Coulter), followed by library preparation using the Ligation library kit (SQK-LSK109, Oxford Nanopore Technologies, ONT) with Native Barcode Expansion (EXP-NBD196, ONT) to allow for multiplexing of libraries. DNA was sequenced on MinION flow cells (R9.4.1, ONT) and basecalled using Guppy version 4.0.14+8d3226e with the dna_r9.4.1_450bps_hac model (ONT).

Illumina-only genome assemblies were generated as reported previously [1]. ONT reads were assembled using Flye version 2.8.1-b1676 [23] (options: --plasmids -g 5250000 --asm-coverage 60) then polished twice with medaka version 1.0.3 [24] using the medaka_consensus module (options: -m r941_min_fast_g303). Finally, assemblies were short-read error corrected using Pilon version 1.23 [25] (two rounds per genome).

Assemblies were annotated using Prokka version 1.14.6 [26], and assembly statistics were generated using the statswrapper.sh module from bbmap version 38.81 [27]. Species, multi-locus sequence types, antimicrobial resistance and virulence determinants were identified using Kleborate version v2.0.1 [6]. Suspected hybrid species were analysed in further detail using GenomePainter version 0.0.8 (https://github.com/scwatts/genome_painter) [28].

### Single nucleotide variant analyses

Single nucleotide variants (SNVs) were identified using RedDog version 1b.10.4 [29]. Isolates were grouped by ST, and the corresponding Illumina reads mapped to the respective completed chromosome and plasmid reference sequences (**Table S1**). As a control, the short-read data from the ST reference isolate was also mapped to its completed assembly. Pairwise SNV distances were calculated for each replicon using the snp2dist script at Figshare (https://doi.org/10.6084/m9.figshare.16609054) [30]. Pairwise chromosomal SNV distances ≤25 were considered indicative of recent strain transmission events, based on previous analyses of *Kp*SC transmission in clinical settings [31–33]. A SNV maximum likelihood phylogenetic tree was also constructed from the SNV alignment using RAxML with a GTR substitution model [30].

### Plasmid analyses

Completed plasmid sequences were analysed using the mob_typer command from Mob-suite version 3.0.0 [34]. The closest matching (best query coverage and identity) plasmid sequences from human-derived *Klebsiella* isolates were identified using NCBI BLASTn search against the public NCBI nucleotide database as of 01/04/21 [35]. A MASH distance tree was generated using mash version 2.1.1 [36], along with R packages ape version 5.4-1 [37] and phangorn version 2.5.5 [38]. The tree was inferred using the FastME algorithm [39] [30].

Visualisations of pairwise plasmid comparisons were generated via EasyFig version 2.2.5 [40] and the BLASTn command, with the following BLAST parameters: Min. length – 1000. Max. e Value – 0.0001, to look at gene content and synteny.

### Genome scale metabolic reconstructions and *in silico* growth predictions

Metabolic diversity of *Kp*SC from flying foxes (n=13 completed genomes only) was explored, in comparison to *Kp*SC from other host and/or environmental sources for which metabolic reconstructions have been published previously (n=37 from [41] and n=22 from [42].

Novel metabolic network reconstructions were generated using cobrapy version 0.20.0 [43] in a conda environment version 4.9.2 [44], running Python 3.6.12 [45]. The *Kp*SC pan metabolic model [41] was used as a reference as previously described [46, 47]. The specific code used to generate models was adapted from [47] and can be found at Figshare (https://doi.org/10.6084/m9.figshare.16609054) [30]. The same approach was performed to update the 22 *K. pneumoniae* reconstructions described in [42]. Model annotations were improved using models iYS1720 [48], iML1515 [49] and iWFL_1372 [50] with a custom script [30]. MEMOTE reports [51] were then generated for each model using the report snapshot option. *In silico* growth capabilities were simulated in minimal media with each of 268 sole carbon, 154 nitrogen, 25 sulfur and 59 phosphorus sources using the flux balance analysis model.optimize method from cobrapy as described previously [41]. All genome accessions, model identifiers are shown in **Table S1**.

To contextualise the metabolic modelling results with regards to population structure, a core genome phylogeny was inferred from a core gene alignment generated with Panaroo version 1.1.2 [52] (options: --mode relaxed -a core --aligner mafft -- core_threshold 0.98 -f 0.90 --merge_paralogs). The resulting alignment of 574,339 variant sites represented 3,319 genes shared by ≥71/72 isolates. RAxML-NG version 1.1.0 [53] was used to infer a maximum likelihood phylogeny with 1000 bootstrap replicates (options: --all --model GTR), best of five runs. RAxML was also used to generate an unscaled, daylight phylogenetic tree in the contexts of plasmid presence [30].

### Data visualisation

Data were visualised using R version 4.0.3 [54] and RStudio version 1.3.1093 [55] with the following software packages: RColorBrewer version 1.1-2 [56], ggplot2 version 3.3.3 [57], reshape2 version 1.1.4 [58], aplot version 0.0.6 [59] and ggnewscale version 0.4.5 [60] [30]. igraph version 1.2.6 [61], ggraph version 2.0.5 [62] and ggforce version 0.3.3 [63] were used for pairwise SNV analyses only. ggtree version 2.4.1 [64] was used for phylogenetic tree visualisation only. pheatmap version 1.0.12 [65], grid version 4.0.3 and gridExtra version 2.3 [66] were used for Flux Balance Analysis visualisation.

## Data availability

These sequence data have been submitted to the Genbank databases under BioProject accession number PRJNA646592. All other data used in this study can be found in supplemental tables and at Figshare (https://doi.org/10.6084/m9.figshare.16609054) [30].

## Acknowledgements

This work was supported, in whole or in part, by the Bill & Melinda Gates Foundation [OPP1175797]. Under the grant conditions of the Foundation, a Creative Commons Attribution 4.0 Generic License has already been assigned to the Author Accepted Manuscript version that might arise from this submission. This work was also supported by Australian Research Council Discovery Project DP200103364.

KLW is supported by the National Health and Medical Research Council of Australia (Investigator Grant APP1176192).

