## Supplementary material for "Transmission of *Klebsiella* strains and plasmids within and between Grey-headed flying fox colonies": Table S5

Table S5: Summary of substrate utilisation for all 72 *Kp*SC isolates in this study based on *in silico* genome scale metabolic models.

|  | **Carbon sources** | **Nitrogen sources** | **Phosphorus sources** | **Sulfur sources** |
| --- | --- | --- | --- | --- |
| **All isolates use** | 135 (49.6%) | 62 (40.0%) | 43 (72.9%) | 10 (40.0%) |
| **Variable usage** | 29 (10.7%) | 5 (3.2%) | 1 (1.7%) | 0 (0.0%) |
| **No isolates use** | 108 (39.7%) | 88 (56.8%) | 15 (25.4%) | 15 (60.0%) |
| **Total** | **272** | **155** | **59** | **25** |
