## Supplemental material document for "Transmission of *Klebsiella* strains and plasmids within and between Grey-headed flying fox colonies"

### Figure S2

Figure S2: Heatmap showing predicted growth capabilities of isolates, clustered hierarchically. Based on *in silico* metabolic modelling in absence of phylogenetic population structure. Species and STs cluster based on metabolic substrate usage, rather than isolation source, consistent with Figure 5.

### Figure S3

Figure S3: GenomePainter image of *K. pneumoniae*-*quasipneumoniae* subsp. *quasipneumoniae* DHQP1002001 species hybrid

### Table S1

Table S1: Isolates used in this study and further genome details, metadata and Kleborate results. See separate excel file.

### Table S2

Table S2: Read mapping statistics generated for SNV calculations within each sequence type and plasmid. See separate excel file.

### Table S3

Table S3: Pairwise SNV calculations generated, used to construct the network graph. See separate excel file.

### Table S4

Table S4: Plasmid table with further details. Closest BLASTn hits are shown, along with Kleborate results. See separate excel file.

### Table S5

Table S5: Summary of substrate utilisation for all 72 *Kp*SC isolates in this study based on *in silico* genome scale metabolic models.

|  | **Carbon sources** | **Nitrogen sources** | **Phosphorus sources** | **Sulfur sources** |
| --- | --- | --- | --- | --- |
| **All isolates use** | 135 (49.6%) | 62 (40.0%) | 43 (72.9%) | 10 (40.0%) |
| **Variable usage** | 29 (10.7%) | 5 (3.2%) | 1 (1.7%) | 0 (0.0%) |
| **No isolates use** | 108 (39.7%) | 88 (56.8%) | 15 (25.4%) | 15 (60.0%) |
| **Total** | **272** | **155** | **59** | **25** |

### Table S6

Table S6: Total metabolic substrate usage profiles for all isolates. Data used for Figure 5. See separate excel file.

### Figshare

Figshare: All scripts and code used to generate data used in this study, along with metabolic models and corresponding genome data can be found at <https://doi.org/10.6084/m9.figshare.16609054>
