## Supplementary figures and images for "Transmission of *Klebsiella* strains and plasmids within and between Grey-headed flying fox colonies"

### Figure S1

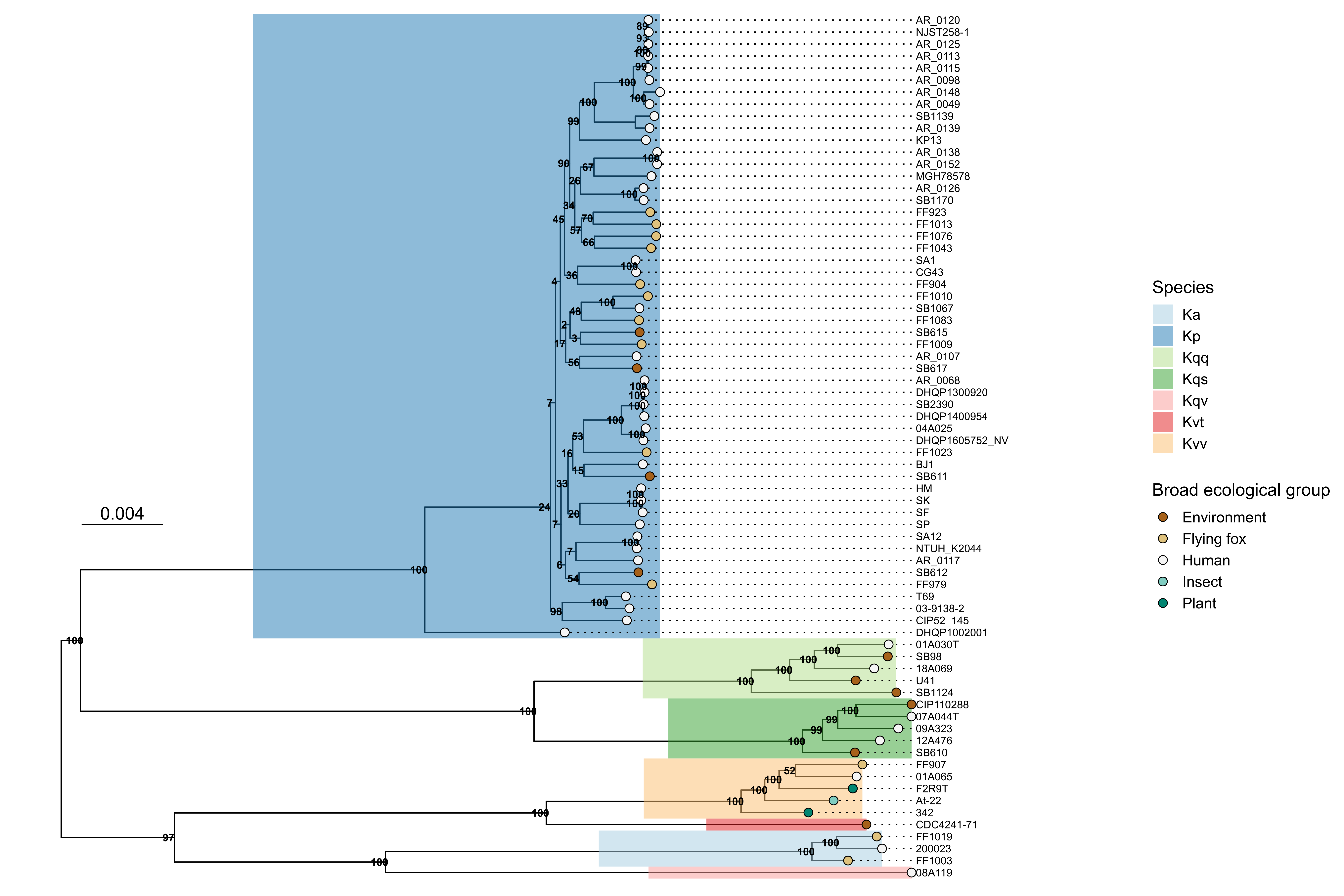

### Figure S2

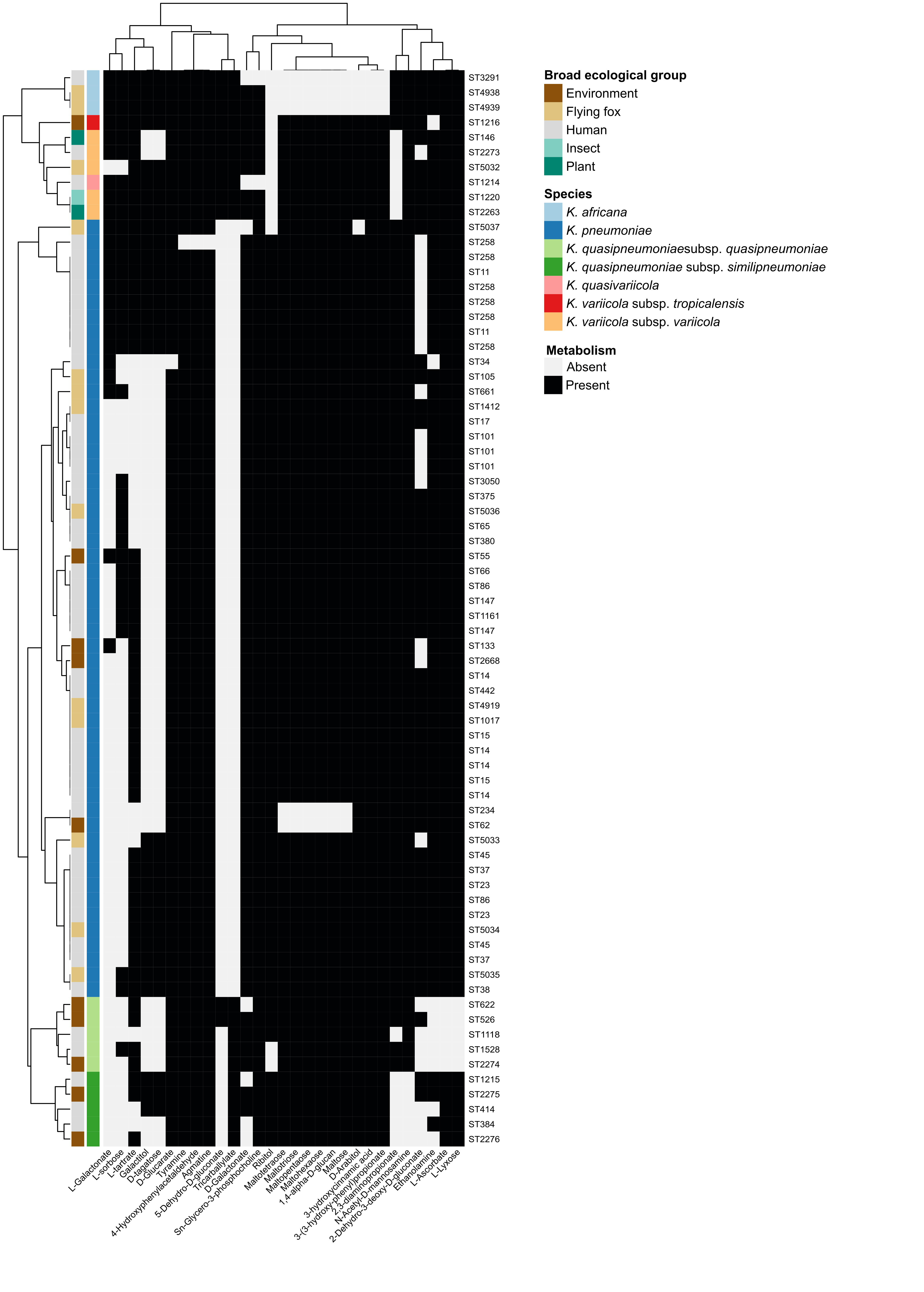

### Figure S3

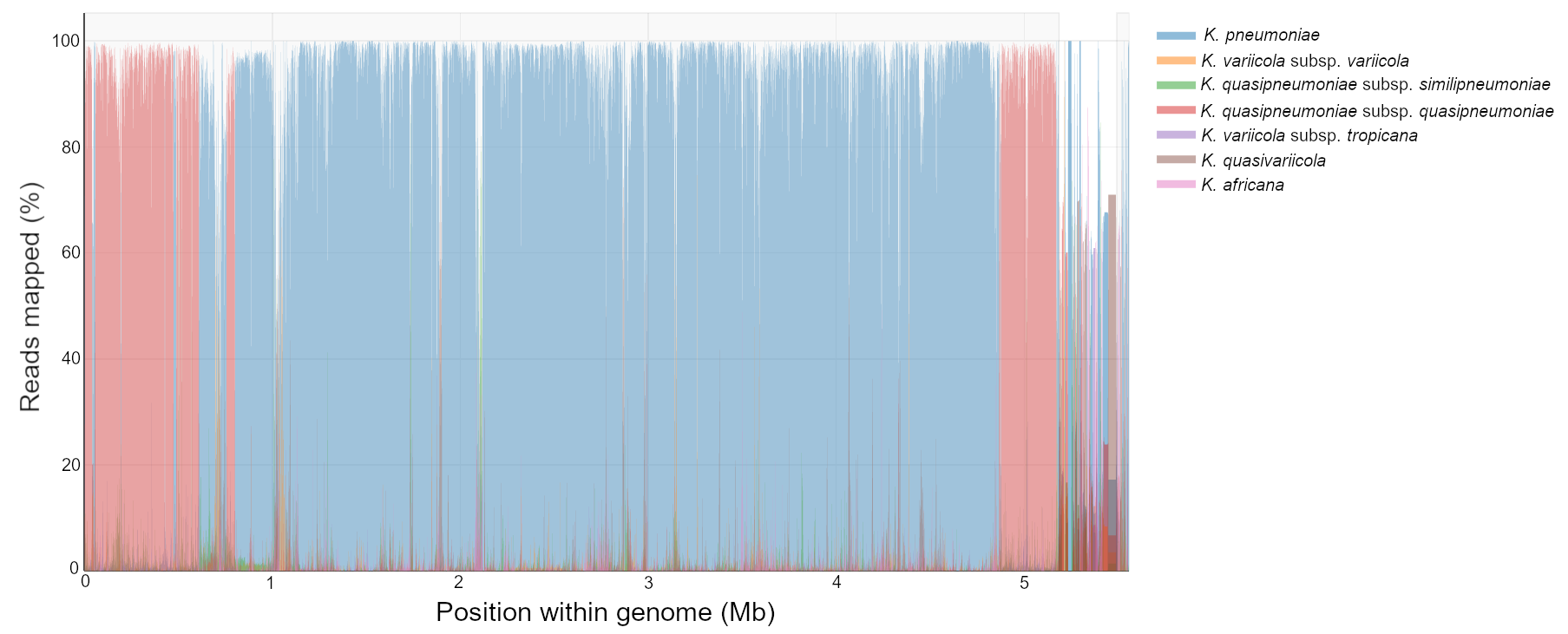
